# ATE1 activates ER-stress and UPR pathways in glioblastoma

**DOI:** 10.1101/2023.09.25.559258

**Authors:** Janaina Macedo-da-Silva, Sueli Mieko Oba-Shinjo, Lívia Rosa-Fernandes, Roseli da Silva Soares, Antonio Marcondes Lerario, Isabele Fattoti Moretti, Talita de Sousa Laurentino, Ricardo Cesar Cintra, Suely Kazue Nagahashi Marie, Giuseppe Palmisano

**Author notes:** these authors jointly directed this work. To whom correspondence should be addressed: Prof. Giuseppe Palmisano, Glycoproteomics Laboratory, Department of Parasitology, ICB, University of São Paulo, Brazil, Av. Prof. Lineu Prestes, 1374, 05508-900 - São Paulo – SP – Brazil.

## Abstract

Post-translational modifications (PTM) have been recognized as a relevant regulation of key processes in cancer pathophysiology, such as cell migration, adhesion, and proliferation. N-terminal protein arginylation is an emerging PTM involved in tumor progression; however, the mechanisms by which this modification influences these events are poorly understood and vary according to cancer type. Glioblastoma (GBM) is an aggressive intra-axial brain tumor associated with poor prognosis, low survival, and high recurrence rate. We performed a study combining *in silico*, *in vitro*, and patients samples analysis to understand the impact of N-terminal protein arginylation in GBM, including overexpression and silencing of *ATE1* in GBM-U87MG cell line with RNASeq analysis, immunofluorescence, and validation of the identified targets at the protein level by immunoblotting. The arginylation pattern differed in GBM compared with non-neoplastic brain tissues, and upregulation of ATE1 was associated with increased tumor cell proliferation. We identified a strong activation of the unfolded protein response (UPR) pathway associated with increased ATE1 level, inducing autophagy and not apoptosis. Protein arginylation in GBM proved to be an important mechanism for tumor growth, with the recycling of cell substrates by autophagy, providing fitness for tumor cells. The expressions of the main markers of UPR and autophagy pathways were validated in human GBM samples, reinforcing the role of ATE1 in the most aggressive brain tumor.

## Introduction

Protein degradation is a well-regulated process to protect cells from misfolded, aggregated abnormal proteins, and some proteolytic pathways can selectively destroy specific subunits of a protein complex acting as protein-remodeling devices. Therefore, this process is involved in many biological functions such as cell cycle, cell differentiation, and responses to stresses. The intracellular degradation is mainly mediated by the ubiquitin-proteasome system (UPS) and by autophagy-lysosome pathways with the participation of molecular chaperones in both systems. Proteins harbor degradation signals, named degrons, which determine the specificity of UPS. Every one of the 20 amino acids in the genetic code can act as a destabilizing N-terminal (Nt)-residue of a N-degron and specific proteolytic systems recognize distinct classes of N-degrons and most often destroy them conditionally. N-degrons are susceptible to enzymatic Nt- modifications, including Nt-arginylation, which involves an enzymatic conjugation of arginine to the Nt by arginyltransferase 1 (ATE1)^1, 2^ . Moreover, ATE1 catalyzes midchain arginylation of Asp and Glu side chains^3^.

Protein arginylation is a post-translational modification (PTM) essential for mammalian embryogenesis and regulation of the cytoskeleton and also influences several cellular functions^4^, including cell survival under stressful conditions as endoplasmic reticulum (ER)-stress^5^, nutrient deprivation^6^, and heat stress^7^.

Cumulative evidence has pointed to the PTM as a relevant process during tumorigenesis and ATE1 expression has been identified as dysregulated in different types of cancer. However, the role of ATE1 in tumor progression is still poorly understood, and its effects differ depending on tumor type. In prostate^8^ and liver cancer^9^, low ATE1 expression was correlated with aggressive tumor features, whereas in melanoma, high ATE1 expression was associated with increased migration, fitness in nutrient deprivation and metastasis, and also with lower survival rate^6^. There is no study about arginylation in brain tumors so far.

Glioblastoma (GBM), a WHO grade 4 astrocytoma, is the most frequent and aggressive intra-axial brain tumor in adults associated with poor prognosis, high recurrence rate, and short overall survival (OS)^10^. Primary and secondary GBM represent distinct tumor subtypes evolving through different genetic pathways^11–13^. Currently, the standard of care for GBM patients consists of surgical resection, followed by radiotherapy and chemotherapy with alkylating drugs such as temozolomide^14^. Several other combinatory therapies have been attempted, but the OS remained around 15-16 months for most patients, and the 5-year survival rate is only 5%^10^. Clinical follow-up of GBM patients has shown that mutations in the *IDH1* gene predict a favorable outcome, with longer OS in those harboring the mutation^15^. Aiming to analyze the effects of arginylation in GBM, we performed *in vitro ATE1* overexpression and silencing in human GBM U87MG cell line, analyzed the involved signaling pathways by RNASeq transcriptomic approach, and validated the selected differentially expressed genes (DEGs) at the protein level in the experimental conditions and also in human GBM samples. We identified a strong activation of the UPR pathway induced by the upregulation of ATE1, leading to increased autophagy and tumor cell survival. Our results corroborated that ATE1 is involved in tumor growth in GBM.

## Methods

### Cell culture

The GBM cell lines T98G and U87MG were obtained from American Type Culture Collection (Manassas, VA, USA) and cultured in Dulbecco’s modified Eagle’s medium (DMEM) (Thermo Fisher Scientific, Carlsbad, CA, USA) supplemented with 10% fetal bovine serum (FBS) (Cultilab, Campinas, Brazil) and antibiotics (100 U/ml penicillin, 100 μg/ml streptomycin). All cells were kept in a humidified atmosphere with 5% CO_2_. Cell line authentication was performed by short tandem repeat DNA analysis with the GenePrint 10 System (Promega, Fitchburg, WI, USA).

### Patients samples

This study was approved by the National Ethics Commission and the local ethics committee under the number 830/01. A written informed consent form was acquired from each patient. All tumor fragments were collected by neurosurgeons of Division of Neurosurgery, Department of Neurology, School of Medicine of University of São Paulo, from 2000 to 2007. Non-neoplastic (NN) brain tissues were collected from epilepsy patients subjected to temporal lobectomy, as previously described^16^. Collected samples were handled following all the necessary biosafety rules for this procedure. All material collected was stored at -80°C. Eight GBM samples and five NN brain tissues were used for protein analysis.

### *ATE1* silencing

The pre-designed small-interfering RNA (siRNA) duplex for the *ATE1* transcript (si*ATE1*) (hs.Ri.ATE1.13.3, 5’-CGGGUGACUUUGCAUUGAUAAAUAA-3’) and non-targeting control (NTC) siRNA were purchased from Integrated DNA Technologies (Coralville, IA, USA). U87MG and T98G cells (2,5 x 10^4^ cells/well) were transfected with Lipofectamine RNAiMAX reagent (Thermo Fischer Scientific) and 10 nmol of si*ATE1* or NTC siRNA in 6-well plates (ATE1-), as recommended by the manufacturer. After incubation for 24h, 48h, and 96h at 37°C and 5% CO_2_.

### CRISPR/Cas9-based transcriptional activation

CRISPR-mediated *ATE1* activation was performed using dCAS9 enzyme (lenti dCas- VP64_Blast, Addgene plasmid #61425), transcription activators (lenti MS2-P65- HSF1_Hygro, Addgene plasmid #61426), and single guide RNA (sgRNA) inserted- plasmide (lenti sgRNA (MS2) zeo backbone, Addgene # 61427). All plasmids were gift from Fen Zhang.

*ATE1* sgRNA guides were cloned into lenti-sgRNA (MS2)_zeo and packaged into lentivirus with Mission Lentiviral Pack Mix (Sigma-Aldrich). Control lentivirus were generated using the empty plasmid to generate control cells (VØ). U87MG cells stably expressing dCas9-VP64-Blast and MS2-P65-Hygro were generated through sequential lentivial transduction and selection with blasticidin (5 μg/mL) and hygromycin (250 μg/mL), respectively, according to protocol previously described^17^. Cells were then transduced with lentiviral containing *ATE1* sgRNA guide (5’- TGGAGAGCGAAGCCTGGCGC-3’) designed in the website http://crispor.tefor.net. Selection was performed with zeocin (250 μg/mL) and then tested for ATE1 expression at transcription and protein levels.

### RNA extraction

Total RNA was extracted from cells using RNeasy Mini Kit (Qiagen, Valencia, CA, USA) following the protocol provided by the manufacturer. The concentration and purity were evaluated using the NanoDrop (Thermo Fisher Scientific), considering values ranging from 1.8 to 2.0 for 260:280 nm absorbance ratio of satisfactory purity. cDNA synthesis was performed from 1000 ng of total RNA using SuperScript III reverse transcriptase, RNase inhibitor (RNaseOUT), random oligonucleotides and oligo(dT), following the recommendations of the manufacturer (Thermo Fisher Scientific). The cDNA obtained was treated with 1 U of RNase H at 37°C for 30 min and 72°C for 10 min to eliminate any hybrids formed. cDNA was then diluted in Tris-EDTA buffer and stored at -20°C for subsequent analysis of genes expressed in these cells. Total RNA was used for library preparation of transcriptome analysis.

### Quantitative reverse transcription real-time PCR (RT-qPCR)

The expression levels of *ATE1* were analyzed by RT-qPCR on the ABI 7500 (Thermo Fisher Scientific). Primers were designed to amplify a region containing 80–120 bp. The primers used were synthesized by Thermo Fisher Scientific as follows: *ATE1* (Forward: CCTTTGCAGTTCACCCTTGG; Reverse: TCTTTCCGTCAAGCCAGTACTG) and hypoxanthine phosphoribosyltransferase (*HPRT*) (Forward: TGAGGATTTGGAAAGGGTGT; Reverse: GAGCACACAGAGGGC-TACAA). Reactions were performed in the final volume was 12 µL per reaction, containing 3 µL of cDNA, 3 µL of primers (final concentration 200 nM) and 6 µL of Power SYBR Green PCR Master Mix (Thermo Fisher Scientific). The amplification conditions included an initial incubation for 2 min at 50°C, 10 min at 95°C, 40 cycles of 15 s at 95°C, and 1 min at 60°C. The expression levels of *ATE1* were normalized to the reference gene *HPRT*. Single product amplification was confirmed by analyzing the dissociation curves. The amplification efficiencies were calculated using serial cDNA dilutions. Assays were performed in triplicates and in two independent experiments.

### Cell Viability

Cell viability assays of U87MG cells controls and silenced or activated for ATE1 expression were performed plating 2.0 × 10^4^ cells per well in a 96-well plate using the PrestoBlue Cell Viability Reagent (Thermo Fisher Scientific). Temozolomide (TMZ) treatment was performed with 0.53mM for 48h. Fluorescence intensity (excitation at 540 nm, emission at 560 nm) was measured using a GloMax-96 Microplate Luminometer (Promega, Madison, WI, USA) on different days. The background consisted of a cell culture medium and was measured for each plate to be used for subtraction from each measurement value. Experiments were performed in octuplicate in two independent analyses. Comparisons between cells with ATE1-modified cells and control cells in each time were analyzed by Two-way Anova followed by post-hoc Bonferroni’s test. The effect of ATE1 silencing was checked in a second GBM cell line by PrestoBlue cell viability assays of T98G cells controls and silenced for ATE1 expression.

### High-throughput RNA sequencing

The transcriptomic profiles of cells were analyzed by RNA-Seq in quadruplicate. Libraries were prepared using the QuantSeq 3′mRNA-Seq Library Prep Kit-FWD by Illumina (Lexogen, Vienna, Austria), using 500 ng of total RNA, following the recommendations of the manufacturer. The library concentration was measured using the Qubit dsDNA HS Assay Kit (Thermo Fisher Scientific), and the size distribution was determined using the Agilent D1000 ScreenTape System on TapeStation 4200 (Agilent Technologies, Carlsbad, CA, USA). Sequencing was performed on the NextSeq 500 platform at the next-generation sequencing facility core SELA at the School of Medicine of the University of São Paulo. Sequencing data were aligned to the GRCh38 version of the human genome with STAR^18^. Downstream processing of the BAM files (merging of different lanes; marking of duplicates) was performed with the bammarkdu-plicates tool from biobambam2; featureCounts was employed to count the number of reads that overlapped each gene^19^. The GFF file containing the gene models was obtained from ftp.ensembl.org. Sequencing quality and alignment metrics were assessed with FastQC and RNASEQC, respectively. Downstream analyses were performed in R using specific

Bioconductor and CRAN tools, and briefly described. Normalization was performed with edgeR using the trimmed-mean method. We used sva to remove occult/unwanted sources of variation from the data. The R-Bioconductor package limma was used to assess differential gene expression in each group, and to perform log2 counts per million reads mapped (CPM) in the transformation of the data. Normalized CPM were converted to z-score for heat map visualization. Principal component analysis (PCA) was performed using the prcomp function from R-stats, and graphically depicted as biplots constructed using ggplot2. Expression data were centered to the mean of each gene. An enrichment map of GO terms was perfomed using the STRING plugin in Cytoscape.

### Immunofluorescence

ATE1 cellular localization in U87MG cell line was analyzed by immunofluorescence. Cells were cultured in monolayer on poly-L-lysine-coated glass coverslips. Cells were fixed with 4% paraformaldehyde, permeabilized with 0.1% Triton X-100 and incubated in blocking solution containing 4% normal goat serum. Subsequently, cells were incubated with the primary antibody anti-ATE1 (1:200, rat, Sigma-Aldrich), anti-calreticulin (CALR) (1:200, rabbit, Abcam) overnight at 37°C, followed by incubation with the anti-rat and anti-rabbit IgG secondary antibody conjugated to Alexa Fluor 488 and 546 (1:400, Thermo Fisher Scientific) overnight at 4°C. Nuclei were stained with 4’,6-diamidino-2-phenylindole (DAPI; Thermo Fisher Scientific). The preparations were analyzed under a confocal microscope Zeiss 510 LSM META and Zeiss 780-NLO (Carl Zeiss Microscopy, Thorn-wood, NY).

### Western blotting

Proteins were extracted from cellular lysates and quantified using the Qubit Protein Assay Kit platform (Invitrogen). A total of 25µg of proteins were separated by SDS-PAGE and electrotransferred to PVDF membranes, which were directly incubated with blocking buffer (5% bovine serum albumin (BSA) in Tris-buffered saline (TBS) at 0.05% Tween- 20 (TBST)) for 1h. Subsequently, samples were incubated with primary antibodies (**Table 1**) overnight and washed three times with TBST. Then, the bands were incubated with the respective secondary antibodies for 1h at room temperature. Immunoreactive bands were detected with the ChemiDoc XRS Imaging System equipment and protein quantification was performed using the ImageJ software. Graphs were plotted using GraphPad Prism version 8.1 software. Bands with statistically significant intensities among groups were evaluated by applying an Ordinary One-way ANOVA, with Tukey post-hoc test (0.05 cut-off).

**Table 1.** Primary and secondary antibodies used in western blotting analyses, with their respective dilutions, reference catalog number, type, and supplier company.

| **Antibody** | **Secondary Antibody** | **Dilution** | **Reference** | **Type** | **Company** |
| --- | --- | --- | --- | --- | --- |
| Anti-ACTB | Goat Anti-Mouse | 1:10000 | #A2228 | Monoclonal | Sigma-Aldrich |
| Anti-AKT | Goat Anti-Rabbit | 1:1000 | #9272 | Polyclonal | Cell Signaling |
| Anti-P-AKT | Goat Anti-Rabbit | 1:1000 | #9271 | Polyclonal | Cell Signaling |
| Anti-ATE1 | Goat Anti-Rat | 1:1000 | MABS436 | Monoclonal | Merck Millipore |
| Anti-ATF4 | Goat Anti-Rabbit | 1:1000 | #11815 | Monoclonal | Cell Signaling |
| Anti-ATF6 | Goat Anti-Rabbit | 1:1000 | #65880 | Monoclonal | Cell Signaling |
| Anti-ATG5 | Goat Anti-Rabbit | 1:1000 | #12994 | Monoclonal | Cell Signaling |
| Anti-BIM | Goat Anti-Rabbit | 1:1000 | #2819 | Monoclonal | Cell Signaling |
| Anti-BIP | Goat Anti-Rabbit | 1:1000 | #3183 | Polyclonal | Cell Signaling |
| Anti-R-BIP | Goat Anti-Rabbit | 1:1000 | ABS2103 | Polyclonal | Merck Millipore |
| Anti-CALR | Goat Anti-Rabbit | 1:1000 | ab2907 | Polyclonal | Abcam |
| Anti-R-CALR | Goat Anti-Rabbit | 1:1000 | ABS1671 | Polyclonal | Merck Millipore |
| Anti-CASP3 | Goat Anti-Rabbit | 1:1000 | #9662 | Polyclonal | Cell Signaling |
| Anti-C-CASP3 | Goat Anti-Rabbit | 1:1000 | #9661 | Polyclonal | Cell Signaling |
| Anti-CYTC | Goat Anti-Mouse | 1:1000 | CTC03 | Monoclonal | Invitrogen |
| Anti-EIF2A | Goat Anti-Rabbit | 1:1000 | #9722 | Monoclonal | Cell Signaling |
| Anti-P-EIF2A | Goat Anti-Rabbit | 1:1000 | #9721 | Monoclonal | Cell Signaling |
| Anti-IRE1A | Goat Anti-Rabbit | 1:1000 | #3294 | Monoclonal | Cell Signaling |
| Anti-P-IRE1A | Goat Anti-Rabbit | 1:1000 | ab48187 | Polyclonal | Abcam |
| Anti-LC3B | Goat Anti-Rabbit | 1:1000 | #2775 | Polyclonal | Cell Signaling |
| Anti-PDI | Goat Anti-Mouse | 1:1000 | #MA3-019 | Monoclonal | Sigma-Aldrich |
| Anti-R-PDI | Goat Anti-Rabbit | 1:1000 | ABS1655 | Polyclonal | Merck Millipore |
| Anti-PERK | Goat Anti-Rabbit | 1:1000 | #3192 | Monoclonal | Cell Signaling |
| Anti-P-PERK | Goat Anti-Rabbit | 1:1000 | #3179 | Monoclonal | Cell Signaling |
| Anti-SQSTM1 | Goat Anti-Mouse | 1:1000 | GT1478 | Monoclonal | Invitrogen |
| Anti-XBP1s | Goat Anti-Rabbit | 1:1000 | #12782 | Monoclonal | Cell Signaling |
| Goat Anti-Mouse | - | 1:4000 | ab6789 | Polyclonal | Abcam |
| Goat Anti-Rabbit | - | 1:4000 | ab6721 | Polyclonal | Abcam |
| Goat Anti-Rat | - | 1:1000 | BA-9400 | Polyclonal | Vector Laboratories |

### Bioinformatics analysis

Public RNA-seq sequencing data from normal brain samples provided by the GTEx project^20^ and from GBM were provided by the TCGA-GBM platform^21^. The packages TCGAbiolinks, TCGAutils, recount, edgeR, and limma were used. The "TCGAquery_recount2" function was used to download data from both platforms, specifying the desired tissue (brain). Data normalization was performed by the "TCGAanalyze_Normalization" function, with the gcContent method. Then, the data were filtered to remove genes with low expression, applying the "TCGAanalyze_Filtering" function^22^. Differently expressed genes were determined using the limma package pipeline^23^. To explore enriched pathways, biological processes, cellular compartments, and molecular functions, the DAVID^24^, g:profiler^25^, and STRING platforms were used, as well as available bioconductor packages such as cluster, factoextra, metboAnalystR, combiroc, pROC, clusterProfiler, flashClust, richplot, ggplot2, and ggstatsplot. Ontologies and pathways were considered enriched if they had a q-value < 0.05 (Benjamini-Hochberg). The Kaplan-Meier curves were built with the survminer and survival packages. Low and high levels of ATE1 was determined based on the average expression.

### Molecular dynamics analysis

The structures corresponding to BIP and R-BIP generated by the AlphaFold^26^ algorithm were minimized in energy using the GROMACS package^27^ after adding appropriate ions to balance the charges. Proteins were simulated using the AMBER99SB^28^ force field and the tip3p water model^29^ in a cubic box. The system was properly equilibrated in the NVT ensemble (constant number of particles, volume, and temperature) for 100ps, followed by pressure equilibration under an NPT ensemble (constant number of particles, pressure, and temperature). The final production of mdrun was performed once at 10ns.

## Results

### The N-degron proteolytic pathway by Nt-arginylation was differentially modulated in normal brain and GBM samples

We analyzed the ATE1 role on proteolytic pathway by arginylation in GBM and validated the main finidngs in *in vitro* assays, as well as in human samples (**Figure S1A**). Innitially, the proteolytic pathway of N-degrons by Nt-arginylation was investigated *in silico* using public RNASeq data provided by TCGA for GBM cases^30^ and GTEx project for normal brain samples^31^. The expression levels of genes related to the five main steps of the N-degron pathway include (1) action of methionine aminopeptidases; (2) N-degron generation at oxidized cysteine residues and deamidation of asparagine and glutamate; (3) formation of the tRNA and ATE1 complex; (4) protein arginylation and (5) proteasomal degradation were compared in GBM (TCGA) and normal brain (GTEx) samples (**Figure 1A**). The principal component analysis (PCA) of these gene expressions showed significant differences in component 1 (44.2%) between tumor and normal samples (**Figure 1B**). Moreover, GBM patients with higher *ATE1* expression levels presented shorter OS by the Kaplan-Meier test (log-rank p=0.047) (**Figure 1C**).

**Figure 1.**
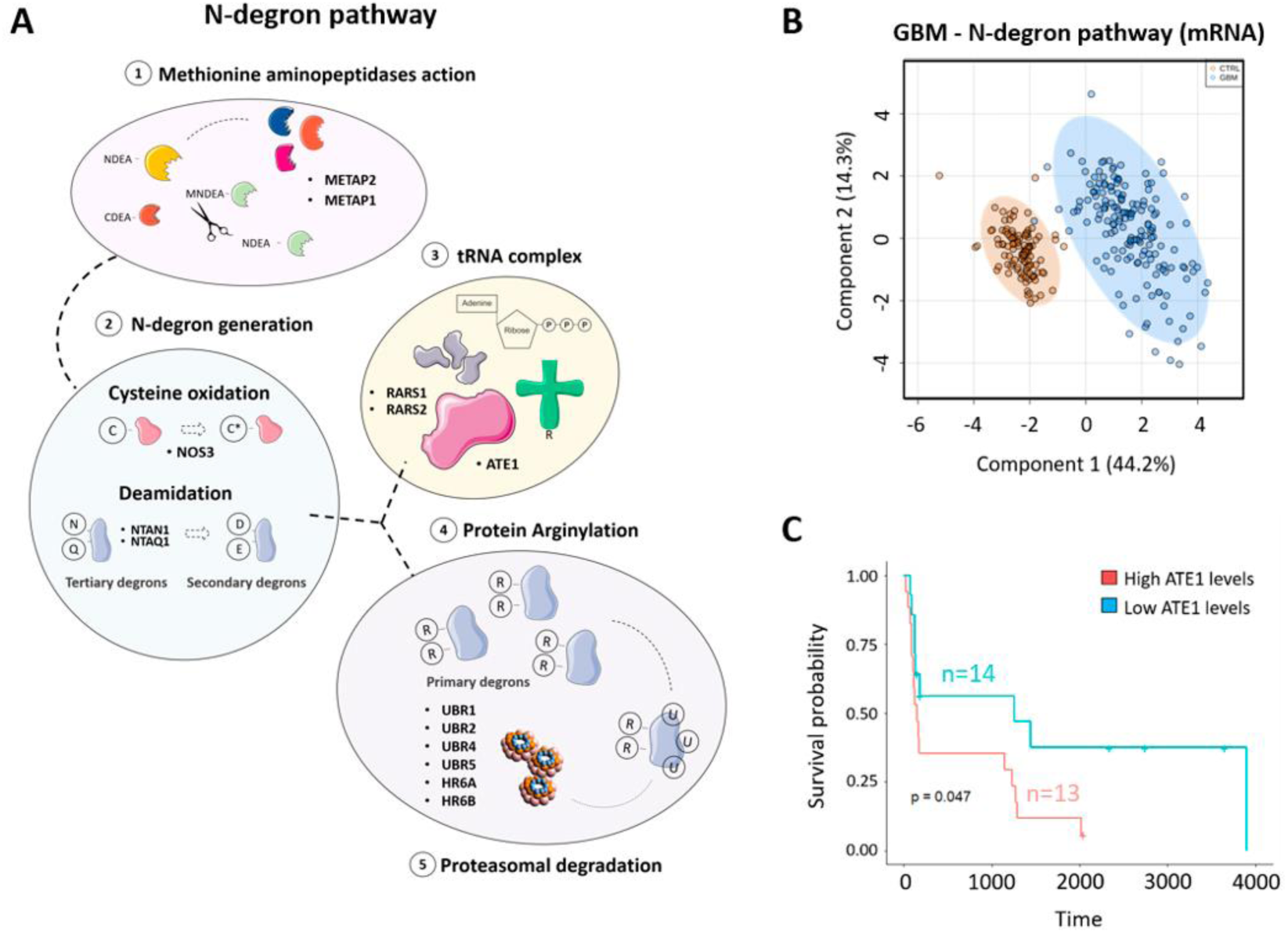
Modulation of ATE1 in glioblastoma (GBM). (**A**) Schematic graph showing the five main steps of the N-degron pathway: (1) Action of methionine aminopeptidases; (2) N-degron generation at oxidized cysteine residues and deamidation of asparagine and glutamate; (3) Formation of the tRNA and ATE1 complex; (4) Protein arginylation and (5) proteasomal degradation. (**B**) Principal component analysis (PCA) was performed using the expression of only the component genes of the N-degron pathway in the dataset provided by The Cancer Genome Atlas Program (TCGA-GBM) and The Genotype-Tissue Expression (GTEx) project. **(C)** Kaplan-Meier curve indicating the probability of survival of patients with high (red, n = 13) and low (blue, n =14) levels of *ATE1*.

### *ATE1* overexpression promoted cell proliferation of U87MG cells

The impact of ATE1 expression level in U87MG cells was addressed by cell proliferation assay in ATE1 overexpressing (ATE1+) and silenced (ATE1-) cells. Up- and down- regulation of ATE1 expression under these conditions was confirmed at gene and protein levels. An increased abundance of arginylated proteins was detected in ATE1+ cells by western blot. In contrast, arginylated protein expressions were significantly decreased in ATE1- cells (**Figure 2A, 2B**). ATE1+ cells showed a significant increase in cell proliferation (p<0.0001 at 24h and 48h,) compared to VØ cells (**Figure 2C**). In contrast, any change of cell of proliferation was observed in *ATE1*-silenced U87MG cells(up to 48h) compared with NTC cells (**Figure 2D**). However, U87MG cells silenced for *ATE1* treated with TMZ presented an increased cell death when compared to non-treated cells (**Figure 2E**).

**Figure 2.**
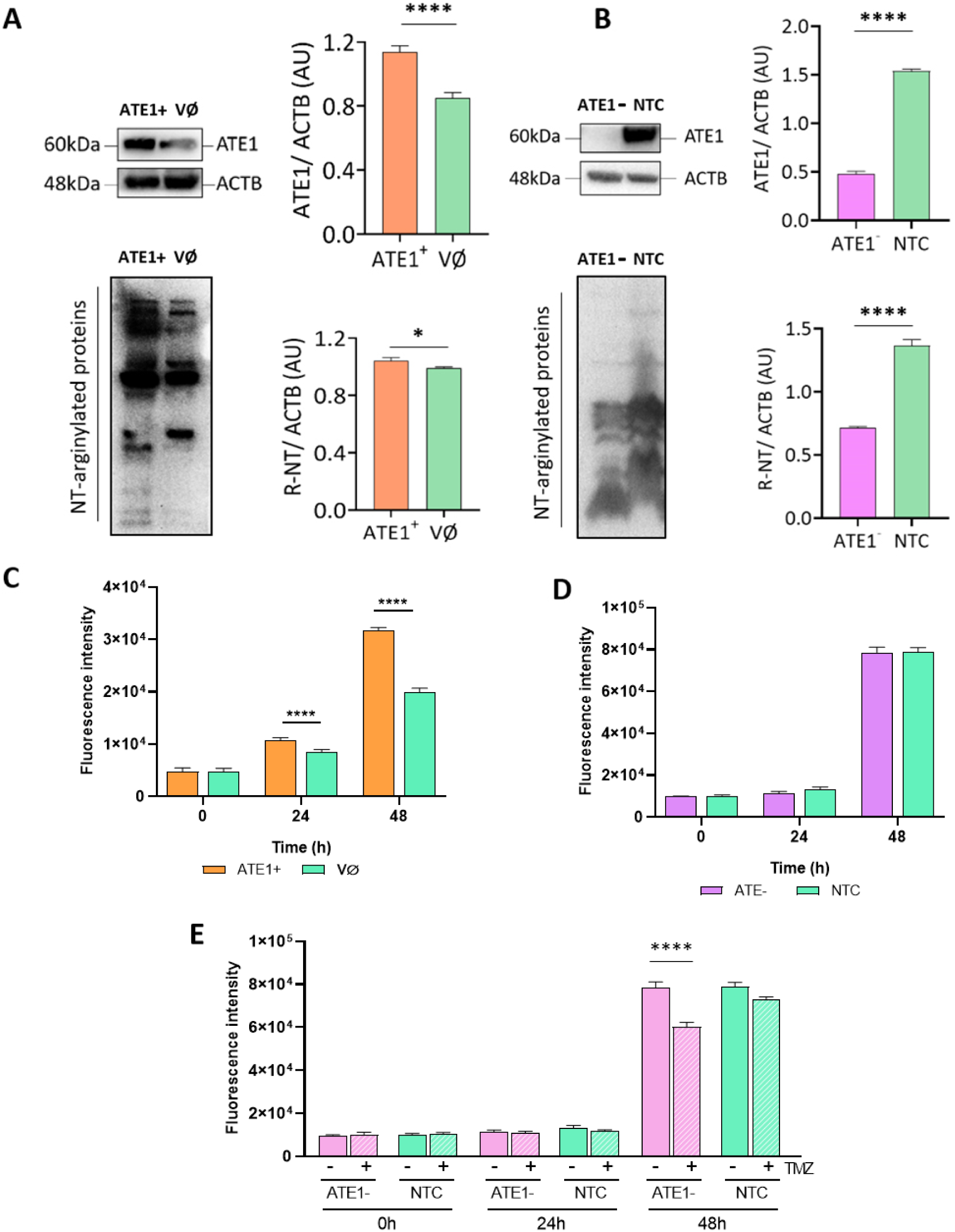
Activation and silencing of ATE1 in GBM-U87MG cell line. (**A**) Representative western blotting images of ATE1 and N-arginylated protein expressions in U87MG cells with overexpressed *ATE1* (ATE1^+^) and control cells (VØ: empty vector). (**B**) Representative western blotting images of ATE1 and N-arginylated protein expressions in U87MG cells with silenced *ATE1* (ATE1^-^) and control (NTC: non-targeting control). (**C**) Growth curves of U87MG ATE1^+^ and VØ cells. (**D**) Growth curves of ATE1^-^ and NTC cells. (**E**) ATE1- cells treated with TMZ (0.53mM) for 48h. **** p<0.0001, * p<0.05 in relation to the control group (Two-way Anova post-hoc Bonferroni’s test).

### U87MG cells overexpressing ATE1 presented an increased cellular response to stress

As a next step, we generated transcriptomic data by RNASeq of ATE1+ cells to analyze the signaling pathways involved in the observed proliferative response. ATE1 upregulation in ATE1+ cells was confirmed at the transcriptional level compared to VØ cells in the RNASeq, with log fold change (FC) = 1, adjPavlue = 3.78e-5 (**Figure 3A**), and the PCA showed clear separation of the gene expression profile between the ATE1+ and VØ cells (**Figure 3B**). The RNASeq analysis yielded a total of 11,994 gene transcripts, including 2,112 DEGs with adjP ≤ 0.05, in ATE1+ compared with VØ cells, being 306 upregulated and 362 downregulated DEGs with log FC ≥ |1|. The downregulated DEGs were enriched for DNA-dependent DNA replication (GO 44786, FDR=0.0049) and for cellular protein complex disassembly (GO 43624, FDR=0.0018). Several genes related to DNA repair and replication (*POLE2*, *GINS1*, *PCNA*); regulation of cell cycle (*CDC7*, *CCNE2*, *GMNN*); maintenance of mitochondrial and nuclear DNA stability (*DNA2*); protein synthesis within the mitochondrion (*MRPL1*); and regulation of microtubule system (*STMN1*) were among the most downregulated DEGs (logFC = -2.4, adjP = 0.0002). Notably, several genes coding for mitochondrial ribosomes (mitoribosomes) (*MRPL*, *MRPS*) were highly connected and significantly downregulated (**Figure 3C, 3E**).

**Figure 3.**
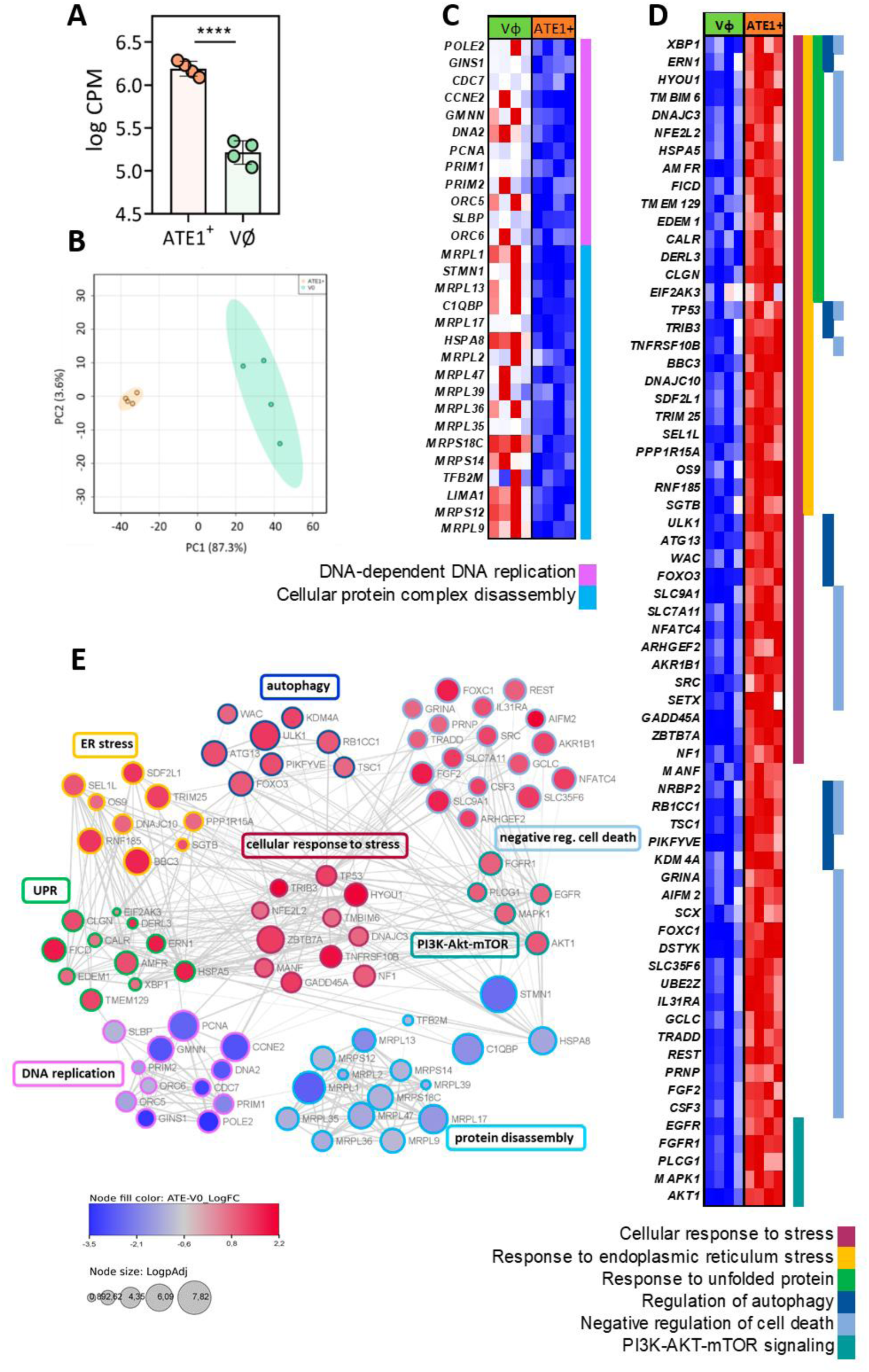
RNA-seq analysis of U87MG cells overexpressing ATE1 (ATE1+). (**A**) Expression of ATE1 in ATE1+ and VØ cells (log CPM), **** p < 0.0001; **(B)** Principal component analysis (PCA) showing the separation between the ATE1+ (orange) and VØ cells (blue); **(C)** Heatmap of downregulated genes comparing ATE1+ and VØ cells grouped in two significant GO enriched pathways **(D)** Heatmap of upregulated genes comparing ATE1+ and VØ cells grouped in six Biological Processes of GO enrichment analysis. The expression values were normalized by z-scores and upregulated genes are presented in red and downregulated genes in blue. **(E)** Connectivity among the up and downregulated genes. The network of proteins identified was analyzed in WebGestalt performed by String app in Cytoscape software. The genes are represented by nodes, the interactions are represented by edges width according to combined score: 0.4 to 0.9. The size of the circles varies according to log adjP value, and the color filling them out varies from red (for upregulated genes) to blue (for downregulated genes) according to logFC (ATE1+ vs VØ).

In contrast, the upregulated DEGs were involved in signaling pathways related to the cellular response to stress (GO 33554, FDR=1.13 x 10^-21^); response to endoplasmic reticulum (ER) stress (GO34976, FDR=2.17 x 10^-28^); response to unfolded protein (GO 6986, FDR=5.33 x 10^-16^); regulation of autophagy (GO 10506, FDR=1.44 x 10^-8^); EGFR tyrosine kinase inhibitor resistance (GO 01521, FDR=0.0224) and negative regulation of cell death (GO 60548, FDR=5.45 x 10^-20^) (**Figure 3D**). Among the 42 upregulated genes in response to cellular stress, 28 genes were related to ER stress, being 16 genes related to unfolded protein, including transcription factors (*XBP1*, *NFE2L2*); ER chaperones (*CALR*, *CLGN*, *DNAJC3, PDI*); members of heat shock protein (Hsp) 70 family (*HYOU1*, *BIP*); Hsp70 binding activity (*FICD*); ER transmembrane receptors related protein degradation (*AMFR*); ER sensor of unfolded proteins (*ERN1*, *EIF2AK3*); misfolded protein binding activity (*EDEM1*); ER degradation of misfolded proteins (*DERL3*); and ubiquitin protein ligase activity (*TMEM129*, *TMBIM6*). Genes coding for proteins related to autophagosome (*ULK*, *ATG5*, *ATG13*) and early endosomes (*PIKFYVE*) were also significantly upregulated in ATE1+ cells compared to VØ cells (logFC = 1.1, adjP < 4.38 x10^-6^). Interestingly, *TP53* was identified as a node interconnecting cellular response to stress, ER stress, autophagy, and negative regulation of cell death (**Figure 3D**).

### ATE1 overexpressed U87MG cells presented an increase in ER stress and unfolded protein response

The transcriptomic analysis of ATE1+ cells demonstrated the upregulation of several genes located in ER, including the ER marker, CALR, that co-localized with ATE1 by immunofluorescence assay, suggesting the presence of ATE1 in ER (**Figure 4A**). The RNASeq data also pointed to upregulation of ER stress, leading to unfolded protein response (UPR); therefore, the identified upregulated targets involved in this pathway were checked at the protein expression level, and counterchecked in ATE1- cells. Interestingly, the Nt-arginylation of BIP was increased in ATE1+ cells and decreased in ATE1- cells. The activation of this signaling pathway occurs when BIP is released when it binds to misfolded proteins accumulated in the ER lumen. Then, the Nt-arginylation of BIP targets the protein for degradation and allows the activation of the UPR signaling pathway (**Figure 4B**).

**Figure 4.**
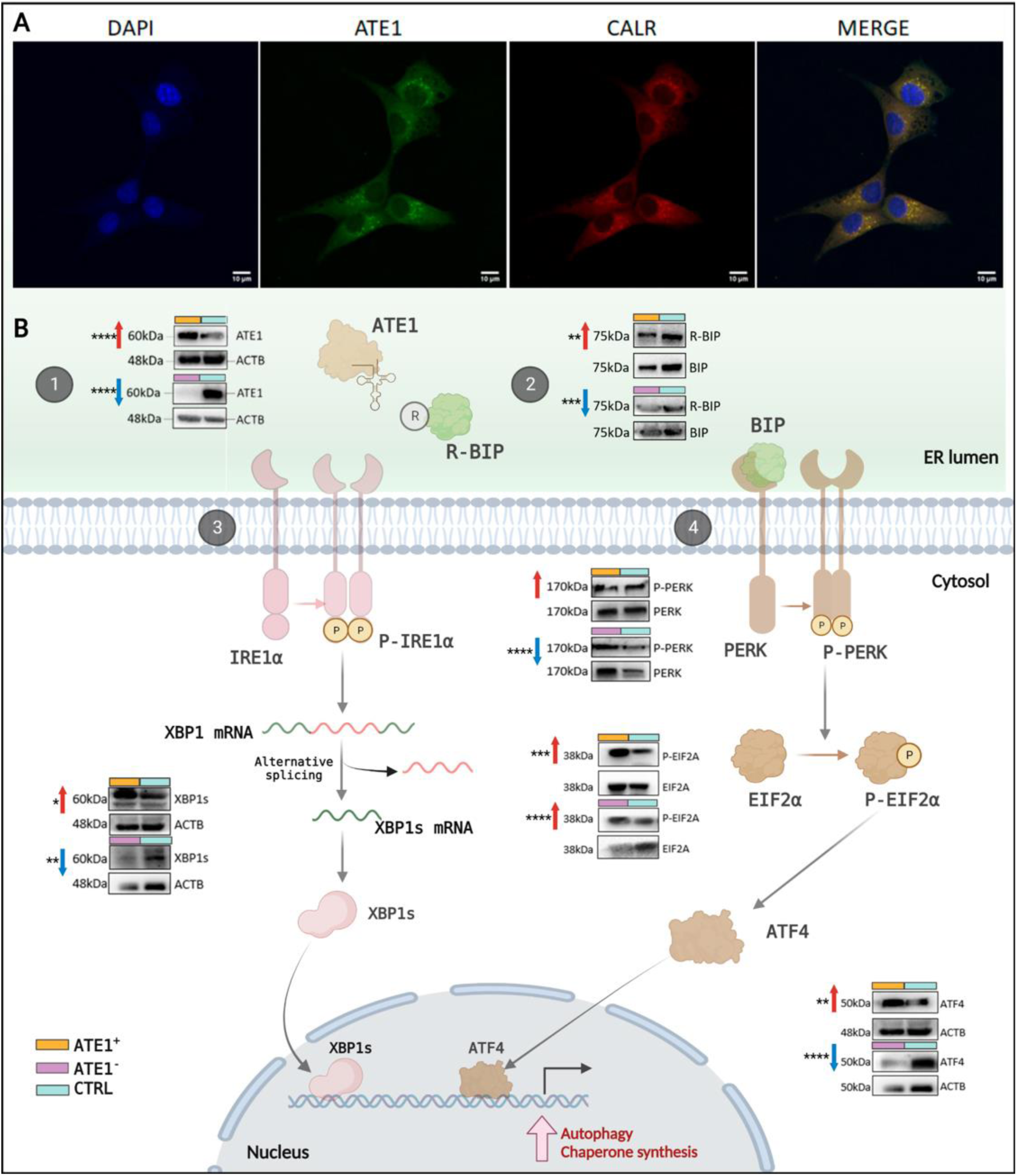
Unfolded protein response (UPR) pathway regulation in U87MG cells. (**A**) Immunofluorescence indicates the co-localization of ATE1 (green) and CALR (red), suggesting the cellular localization of ATE1 in the endoplasmic reticulum (ER). **(B)** Representative western blotting analysis of UPR pathway in activated (ATE1+) and silenced (ATE1-) U87MG cells. Up red arrows indicate upregulation and down blue arrows indicate downregulation of activated or silenced cells relative to controls. **** p <0.0001; *** p<0.001; **p<0.005; * p< 0.05 in relation to the respective control groups for triplicate analysis. ATE1 (1) promotes BIP arginylation (2), which releases IRE1α (3) and PERK (4) for downstream activation leading to the upregulation of XBP1 and ATF4 transcription factors with essential roles in autophagy. XBP1 undergoes unconventional splicing giving raise to a spliced form (XBP1s) that triggers autophagy. Created with BioRender.

In fact, molecular dynamics (MD) analysis showed that the addition of Nt arginine to BIP significantly altered the structure of the protein (**Figure S3**). Comparison of the mean square deviation (RMSD) distribution showed differences between the backbone of the structures, which become noticeable at 10ns of simulation (p-value < 0.001). The flexibility calculated by the root-mean-square-fluctuation (RMSF) showed more flexible residues in the structure of BIP compared to R-BIP. The Nucleotide-binding (NBD) and Substrate-binding (SBD) domains presented greater flexibility and better-defined peaks, revealing greater movement at 10ns. The Ramachandran plot showed a displacement of the psi and phi angles of the beta-sheet and right alpha-helix structures (**Figure S3**) of BIP in relation to the arginylated form.

In agreement, the ER-resident transmembrane detector of unfolded protein: (PKR)-like ER kinase (PERK, coded by *EIF2AK3*) was activated by phosphorylation in ATE1+ cells compared to VØ cells, and a significantly lower protein acttivation was observed in the ATE1- cells. Likewise, the activation of the downstream effectors EIF2A and the upregulation of transcription factor ATF4 were also observed in ATE1+ cells, in contrast of ATF4 downregulation in ATE1- cells (**Figure 4B**). The activation of the UPR branch of Inositol-requiring enzyme 1 (IRE1, coded by *ERN1*) was confirmed by upregulation of spliced form of the downstream transcription factor (sXBP1) in ATE1+ cells, with corresponding downregulation in ATE1- cells (**Figure 4B**).

### ATE1 levels altered the regulation of apoptosis, autophagy, chaperone synthesis, and proliferation pathways

Activation of the UPR pathway may stimulate the transcription of genes involved in apoptosis, autophagy, and chaperone synthesis to restore cellular homeostasis. Looking for the intrinsic pathway of apoptosis, the pro-apoptotic protein BIM, the Bcl-2 interacting mediator of cell death, was analyzed regarding its three isoforms. The isoform BIM_L_, which initiates apoptosis by Bax activation, was downregulated in both ATE1+ and ATE1- cells. In contrast, the isoforms BIM_EL_ and BIM_S_ were upregulated. Additionally, cytochome C (CYTC) protein presented higher levels in ATE1+ cells and lower levels in ATE1- cells compared to respective control cells. Despite these upstream signaling that supports the apoptosic process, we observed reduced levels of cleaved caspase 3, one of the best known marker of apoptosis, in both cellular models (ATE1+ and ATE1-) (**Figure 5**), indicating that apoptosis was not activated. Such result was convergent to no difference observed in cell death assay by flow cytometry in both conditions (data not shown). Accordingly, transcriptome analysis evidenced an upregulation of genes that regulated negatively cell death. Altogether, the alteration of ATE1 expression did not impact apoptosis. In contrast, RNASeq data pointed to the induction of cell proliferation through the activation of the EGFR/PI3K/Akt axis, which was confirmed at the protein level with a considerable increase in phosphorylated AKT (P-AKT) level in ATE1+ cells, and an equivalent reduction of P-AKT level in ATE1- cells (**Figure 5**).

**Figure 5.**
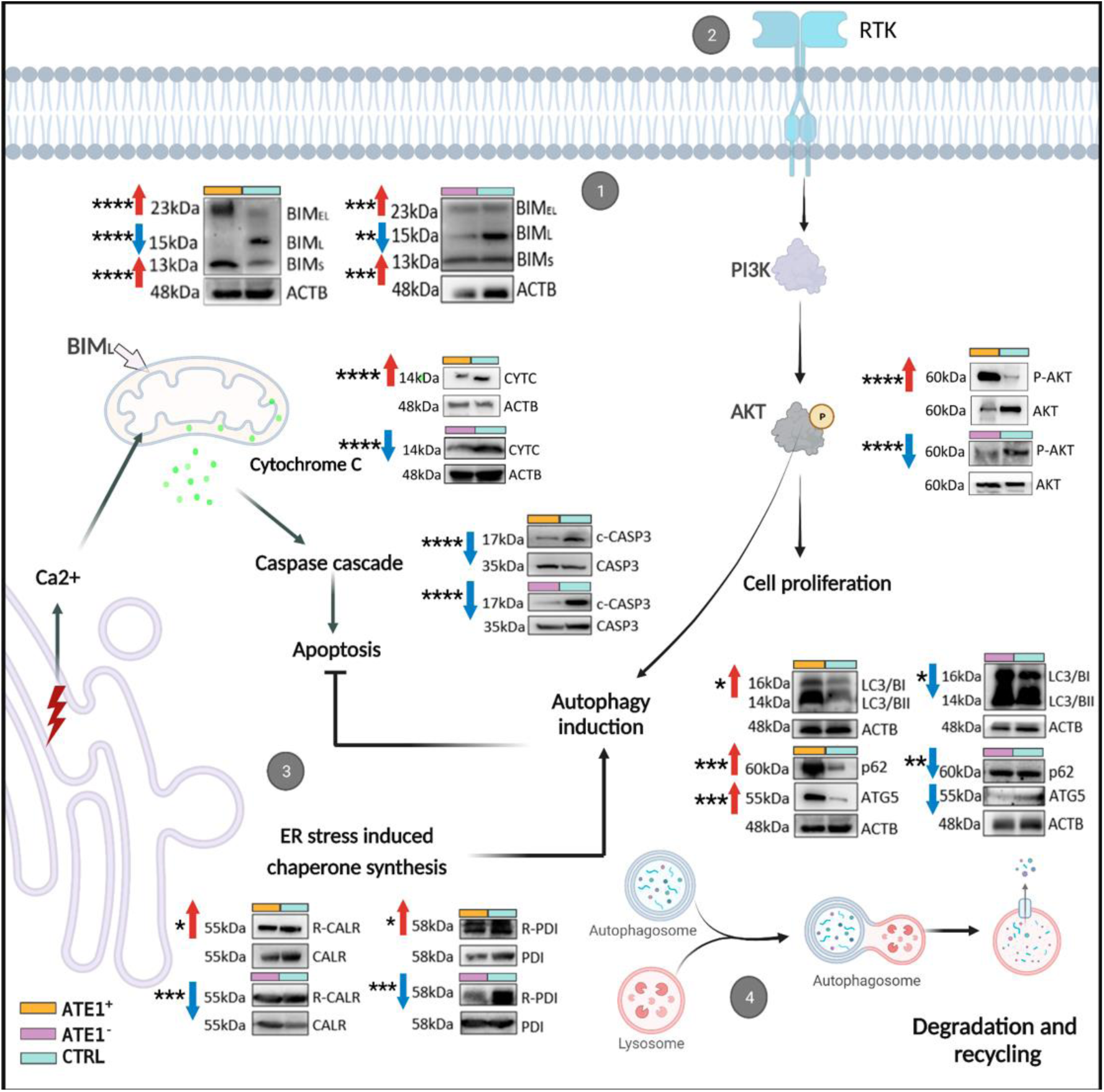
Regulation of apoptosis, RTK-induced PI3K/AKT signaling pathway, ER stress-induced chaperone synthesis, and autophagy in U87MG cells. Representative western blotting analysis of BIM, CYTC, CASP3, AKT, p62/SQSTM1, ATG5, LC3B, R-CALR and R-PDI proteins in ATE1+ and ATE1- U87MG cells. Up red arrows indicate upregulation and down blue arrows indicate downregulation of activated or silenced cells relative to respective controls. **** p <0.0001; *** p<0.001; **p<0.005; * p< 0.05 in triplicate analysis. (1) BIM isoforms (BIMEL, BIML, and BIMs promote apoptosis, being BIML the most potent inducer by associating with BAX activation in mitochondria with cytochrome C (CYTC) release, and cleavage of caspase 3, the final executor of apoptosis. (2) Receptor tyrosine kinase (RTK) signaling pathway by PI3K/AKT promotes cell proliferation and induction of autophagy. (3) ER stress induces chaperone synthesis as calreticulin (CALR) and protein disulfide isomerase (PDI). Both are arginylated (R-CALR, R-PDI). (4) Autophagy with LC3, p62, and ATG5 in autophagosomal formation, cargo collection, degradation, and recycling of cellular substrates for the reprogramming of proteolytic flux. Created with BioRender.

Additionally, RNASeq data pointed to an increase in autophagosome formation and autophagy in ATE1+ cells. ATG5, essential for autophagic vesicle formation, p62/SQSTM1, and ratio of LC3B proteins (LC3/BII and LC3/BI) related to autophagosome formation, were significantly upregulated in ATE1+ cells and downregulated in ATE1- cells, respectively, indicating activation of autophagy in ATE1+ cells (**Figure 5**).

Induction of transcription of chaperone-encoding genes is one of the downstream effects of UPR pathway activation. Two chaperones susceptible to Nt-arginylation, CALR and PDI, which respectively aid in the proper folding of glycoproteins and peptide disulfide chains, were arginylated in ATE1+ cells and downregulated in ATE1- cells, suggesting that protein-folding homeostasis is not restored in ATE1+ cells.

Considering the heterogeneity of GBM, the effect of ATE1 silencing was analazed in a second GBM cell line, T98G. A reduction of ATE1 and Nt-arginylated proteins was also observed in this cell line by western blot. Interestingly, although a decrease of R-BIP protein was observed in T98G ATE1- cells, in contrast to U87MG ATE1- cells, XBP1s was not decreased. Instead, a downregulation of P-EIF2A was observed (**Figure S2B**), indicating slightly distinct downstream UPR alteration in the T98G cell line. Moreover, p62 was also increased T98G ATE1- cells, indicating an increment fo autophagy, which was convergent with an increase of T98G cell proliferation in 48h compared with NTC cells (p<0.001, two-way Anova, post hoc Bonferronís test) (**Figure S2B**).

### Increased ATE1 protein levels and UPR activation were confirmed in human GBM samples

Human GBM samples and non-neoplastic brain samples were analyzed by immunoblotting of the targets differentially expressed to validate the *in vitro* findings. A significant increase of ATE1 was confirmed in GBM samples. Notably, an increased abundance of proteins related to UPR response, R-BIP, PERK, ATF4, XBP1, and a significant increase of ATG5 related to autophagy were observed in GBM samples compared to non-neoplastic control samples (**Figure 6**).

**Figure 6.**
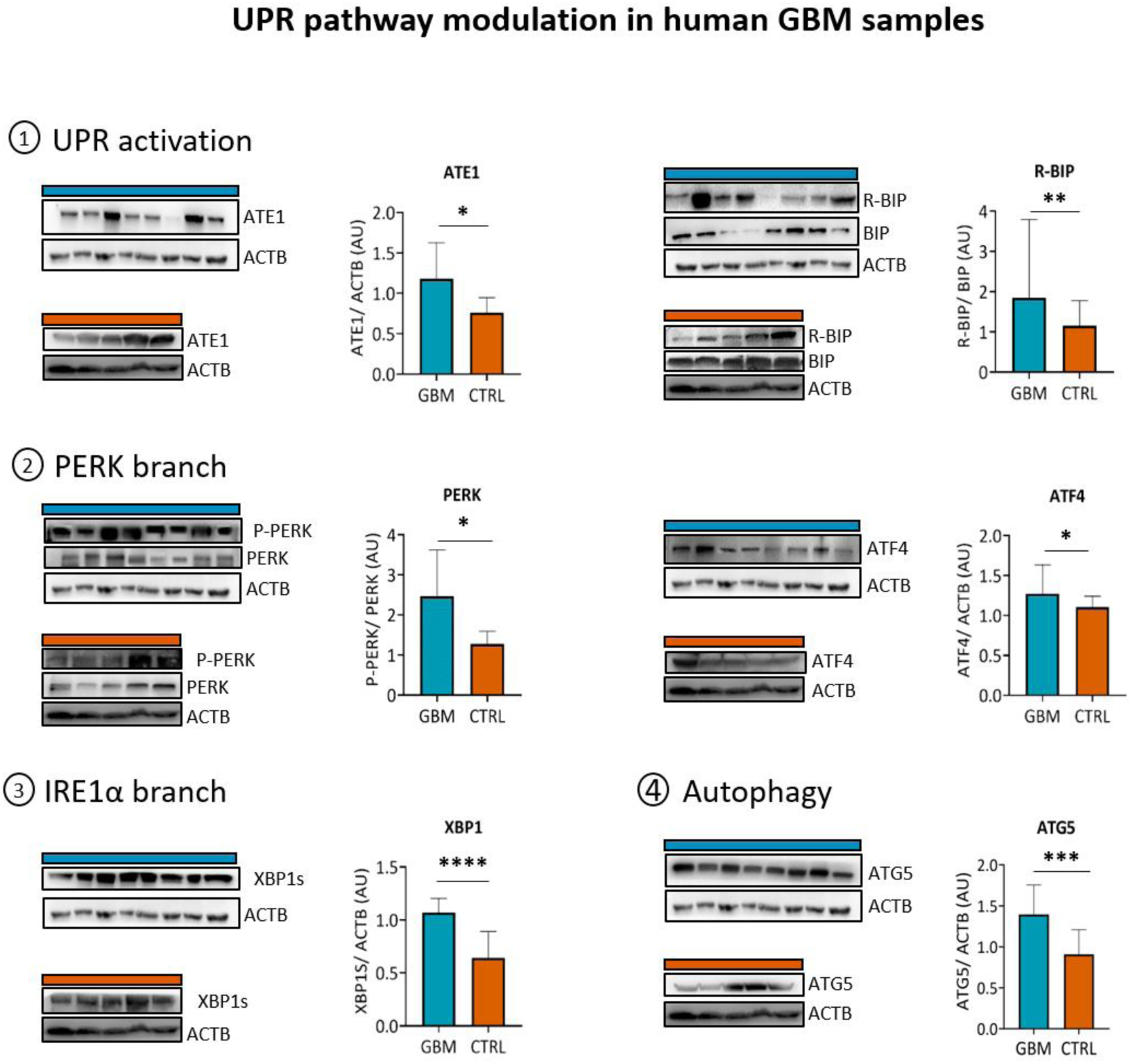
Regulation of ATE1 and UPR proteins in human GBM and non-neoplastic brain tissues. Western blotting indicating modulation of ATE1, proteins of the UPR pathway (PERK, BIP, XBP1, and ATF4) and autophagy (ATG5) in tumor (n=8) and non-neoplastic brain (n=5) tissue samples. **** p <0.0001; *** p<0.001; **p<0.005; * p< 0.05 in relation to the control group.

## Discussion

In this study, we addressed the changes triggered by expression alteration of ATE1, the enzyme that conjugates arginine to Nt, a post-translational modification named arginylation, in the GBM-U87MG cell line and validated the *in vitro* findings in human GBM samples. The preliminary *in silico* analysis of the TCGA-RNASeq dataset for GBM cases^30^ and GTEx for normal brain samples^31^ demonstrated a differential activation of arginylation between GBM and normal brain samples, and a poorer clinical outcome in GBM cases with increased ATE1 expression. Increased ATE1 level has already been correlated with increased cell migration and metastatic events in melanoma^6^. Although the mechanisms of tumorigenesis related to arginylation are not completely elucidated, arginylation has been linked to cell proliferation through AKT^32^. Our analysis of RNASeq of U87MG-ATE1+ cells showed other pathways of interest in cancer, mainly ER stress, UPR response, and autophagy.

UPR activation may occur in different conditions, including hypoxia, oxidative stress, nutrient deprivation, and acidosis^33, 34^. Moreover, protein arginylation has been described as regulating cellular stress response^35–37^. Here, we demonstrated that upregulation of ATE1 was associated with UPR response that triggers cell survival in U87MG cells, as previously described in melanoma cells, where increased cell survival was related to ATE1 overexpression under stress condition triggered by serum deprivation^6^. In particular, we observed increased BIP arginylation in U87MG-ATE1+cells (**Figure 4**) and human GBM samples (**Figure 6**). The BIP protein is fundamental in the activation of the UPR pathway, as it binds to sensors of the UPR, PERK, and IRE1α, and maintains the downstream signaling cascade inactive^38^. The arginylated BIP may also migrate into the cell cytosol and be selectively delivered to autophagic vacuoles by binding with p62^39^. We hypothesized that increased BIP arginylation may activate UPR through the release of PERK and IRE1α sensors for downstream cascades activation, resulting in upregulation of ATF4, a transcriptional master regulator of amino acid metabolism and stress responses^40^, and XBP1, a key transcription factor for cell fate determination in response to ER stress^41^. XBP1 mRNA undergoes unconventional splicing through IRE1α in response to ER stress, giving rise to a 56kDa spliced form (XBP1s) with intact transcriptional activity. XBP1s may trigger autophagic response through transcriptional regulation of Beclin-1 in endothelial cells^42^. Moreover, R-BIP may bind upregulated p62 and induce autophagy (**Figure 5**), favoring tumor cell survival. In fact, previous reports have shown tumor cell survival associated with UPR activation as an adaptive mechanism during disease progression^43–46^. UPR activation was associated with malignant progression and poor prognosis in prostate cancer^43^, and the activation of PERK cascade promoted mouse embryonic fibroblasts adaptation and angiogenesis in response to hypoxic stress^46^. In GBM, the activation of IRE1α cascade resulted in tumor growth and angiogenesis^47–49^, and PERK cascade was associated with cell viability and control of survival rate under stressful conditions as low glucose in U87MG and U251 GBM cell lines^50^. Additionally, RNA expression profile analysis of normal brain tissue and GBM cell lines revealed increased ATF4 and chaperone expressions in response to ER stress and oxidative stress^51^.

An important downstream effect of the UPR is to stimulate apoptosis by caspase activation^52^. However, we did not find an increase in apoptosis (**Figure 5**) in U87MG- ATE1+ cells. On the other hand, we observed different expression levels of BIM isoforms. BIM, the Bcl-2 interacting mediator of cell death, is a member of the BH3-only family of pro-apoptotic proteins. BIM is bound to the dynein light chain (LC8) and localizes to the mitochondria following pro-apoptotic stimulus to start the mitochondrial cell death pathway. BIM alternative splicing yields three major isoforms: BIM_EL_, BIM_L_, and BIM_S_. In particular, BIM_L_ can directly neutralize Bcl-xL to promote apoptosis by Bax activation^53^. BIM_L_ has also been related to cell maintenance or autophagic activities, through its interaction with dynein which facilitates the loading, fusion, and positioning of lysosomes^54^. Moreover, a previous study demonstrated that pro-apoptotic protein fragments Asp-Bcl-xL, Arg-Bid, and Arg-BIM_EL_ are short-lived substrates by selective degradation of the N-degron pathway, which contributes to preventing the pro-apoptotic signal from reaching the point of commitment of apoptosis^55^. Therefore, upregulation of ATE1 may lead to an anti-apoptotic action, which was convergent to the reduced expression of the cleaved caspase 3, the final executor of apoptosis in U87MG-ATE1+ cells (**Figure 5**).

Notably, we also observed an increase in autophagy markers (LC3, p62, ATG5) under ATE1 activation (**Figure 5**). LC3 is an autophagosome marker^56^; p62/SQSTM1/A170 is an autophagy-adaptor protein with an important role in directing and compacting ubiquitinated cargos towards autophagy^57^; and ATG5 is considered to be essential for the induction of autophagy^58^. During stressful conditions, autophagy is critical for the maintenance of cellular homeostasis^59^, regulating the pro-growth signaling cascade and activating the metabolic rewiring of tumor cells to support their growth even in nutrient- deprived microenvironments^60, 61^. Then, when the proteostasis by the UPS is perturbed, Nt-Arg redirects the cellular wastes to macroautophagy through p62 self-polymerization upon its binding to the Nt-Arg, which facilitates cargo collection and lysosomal degradation of p62 cargo-complexes. Therefore, the Nt-Arg arginylated proteins contribute to reprogramming global proteolytic flux under stresses^12^. The importance of autophagy in GBM is not fully understood yet, though previous reports indicated its role in cancer progression and its association with poor survival in GBM patients^62–66^.

Interestingly, the *in vitro* findings in U87MG-ATE1+ cells recapitulated the characteristics of human GBM samples, which showed an increased ATE1 expression with an increment of proteins related to UPR response, R-BIP, PERK, ATF4, XBP1s, and a significant increase of ATG5, corroborating the association of ATE1 increase with induction of autophagy.

Interestingly, upregulation of ATE1 also induced increased arginylation of CALR (R-CALR) (**Figure 5**), an ER-resident protein that operates as a chaperone and calcium buffer to assist correct protein folding within ER^67^. CALR is also implicated in the reduction of apoptosis, and its arginylated form is recruited to stress granules to promote not only its deactivation but also the deactivation of pro-apoptotic proteins, interrupting the apoptosis signaling cascade^68, 69^. PDI, another chaperone located and functioning in ER, was also arginylated and upregulated in U87MG-ATE1+ cells. PDI plays a significant role in protein folding and quality control in the calcium-rich oxidative environment of the ER^70^. The immunofluorescence assay showed ATE1 co-stained with CALR, suggesting the ATE1 localization in ER. Although this finding seemed reasonable, further studies are needed to confirm this observation mechanistically.

Downregulated genes in U87MG-ATE1+ cells were associated to mitochondrial ribosomal proteins (MRPs), essential elements in mitochondrial translation, that have been implicated in biogenesis, metabolism, and apoptosis^71^. They can also modify gene expression, and genomic stability^72^, delay cell cycle progression, and induce apoptosis by regulation of p21WAF1/CIP1, p27Kip1, and p53^73^. MRPL41 is a pro-apoptotic factor that interacts with BCL-2; MRPS29 is associated with Fas receptor-linked death-inducing signaling, and MRPS30 is associated with programmed cell death^74^. The downregulation of the MRP family reinforces the decrease in the activation of the apoptosis cascade identified in U87MG ATE1+ cells. Further analysis of the observed MRP signature in the context of arginylation would be worthwhile in future studies.

In summary, based in the present GBM cohort, corroborated in an extended public GBM dataset, the arginylation proved to be an important mechanism for GBM tumor growth. Activation of the UPR pathway mediated by increased ATE1 level may directly affect the tumor phenotype, with increased recycling of cell substrates by autophagy incrementing the tumor cell fitness. Considering the high heterogeneity of GBM, the specific downstream activation of UPR pathway may vary and need to be determined to allow efficient targeting for therapeutical purposes.

## Acknowledgments

We are grateful for the financial support provided by the São Paulo Research Foundation (FAPESP, grants processes n° 2018/18257-1 (GP), 2018/15549-1 (GP), 2020/04923-0 (GP), 2021/00140-3 (JMDS), 2021/01207-4 (TSL) 2020/02988-7 (SKNM, SMOS); 2015/26722-8 (CW), 2020/12277-0 (EES), 2020/06409-1 (ELD); by the Conselho Nacional de Desenvolvimento Científico e Tecnológico (“Bolsa de Produtividade” (SMOS, SKNM and GP); by Fundação faculdade de Medicina (FFM-SKNM); by the Coordenação de Aperfeiçoamento de Pessoal de Nível Superior (CAPES bolsa PNPD to LRF, #88887.600107/2021-00 to IFM)).

