## Supplementary figures and images for "ATE1 activates ER-stress and UPR pathways in glioblastoma"

### Figure S1

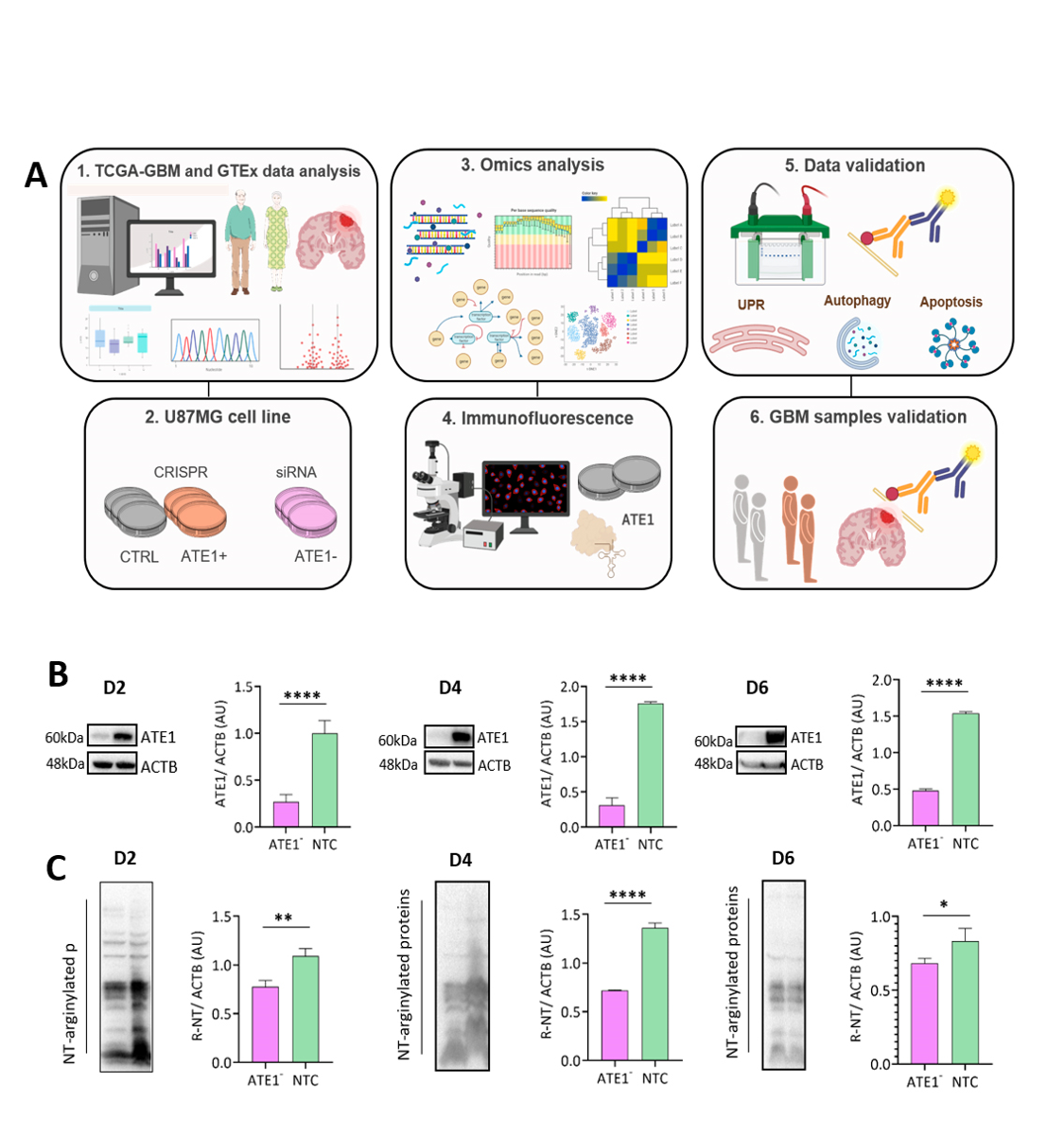

### Figure S2

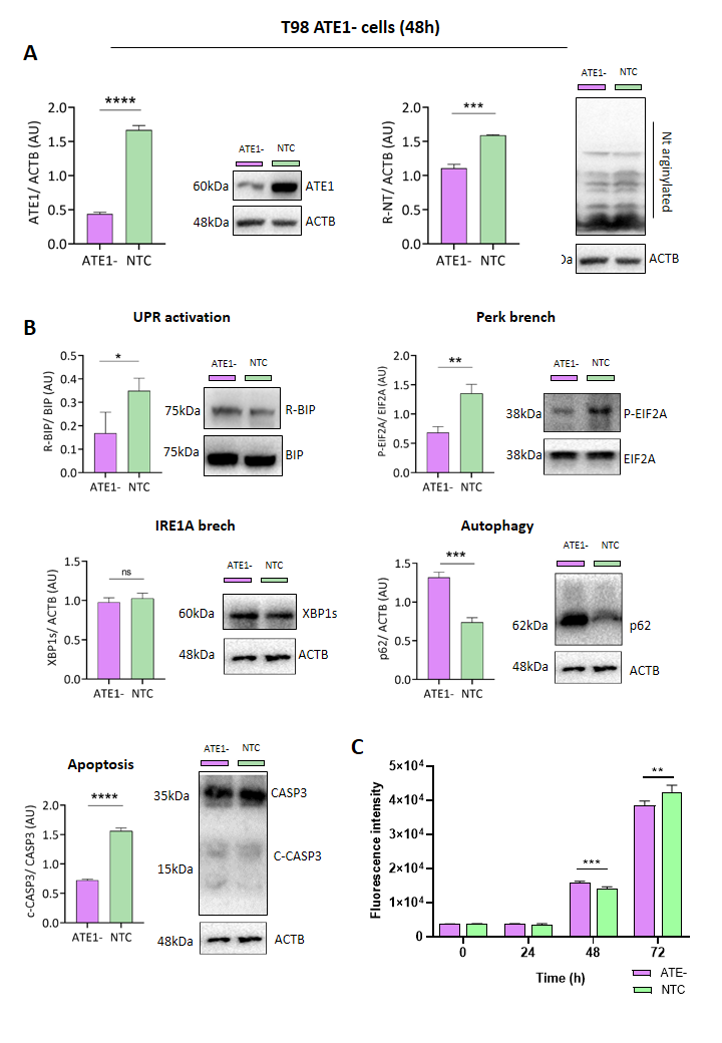

### Figure S3

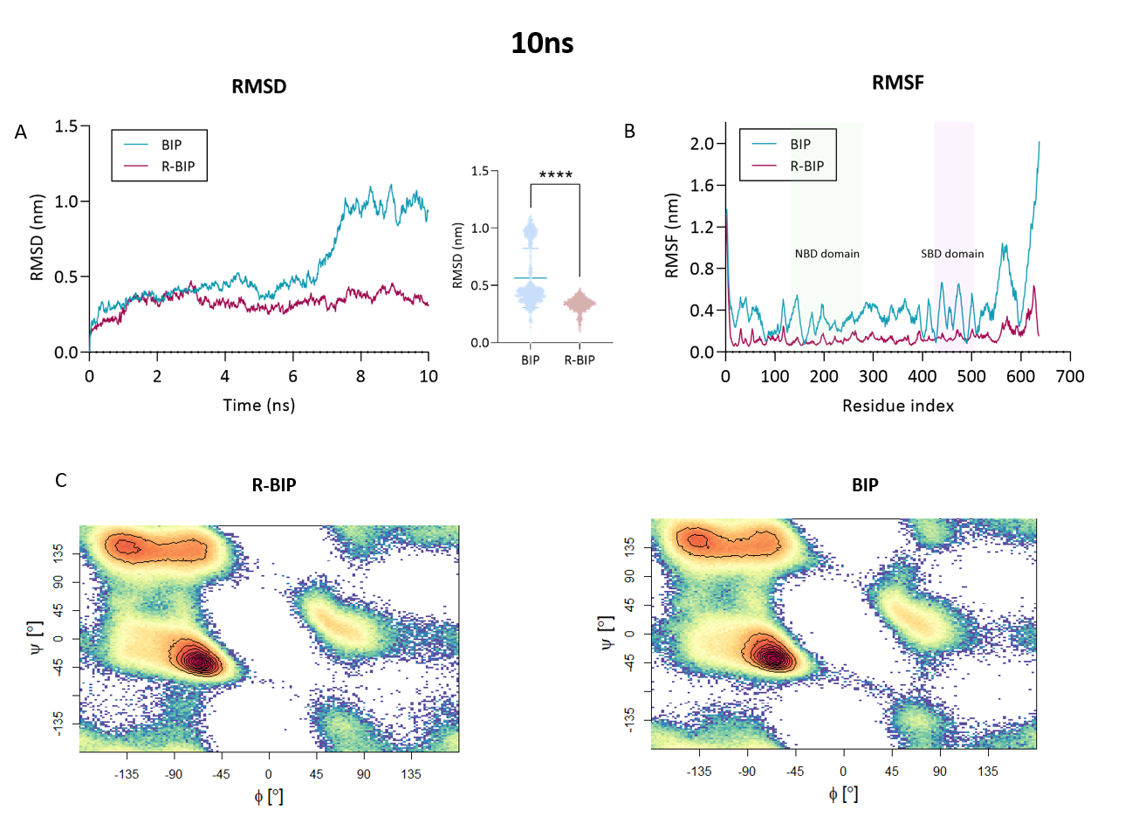
